# Spatial domain detection using contrastive self-supervised learning for spatial multi-omics technologies

**DOI:** 10.1101/2024.02.02.578662

**Authors:** Jianing Yao, Jinglun Yu, Brian Caffo, Stephanie C. Page, Keri Martinowich, Stephanie C. Hicks

## Abstract

Recent advances in spatially-resolved single-omics and multi-omics technologies have led to the emergence of computational tools to detect or predict spatial domains. Additionally, histological images and immunofluorescence (IF) staining of proteins and cell types provide multiple perspectives and a more complete understanding of tissue architecture. Here, we introduce Proust, a scalable tool to predict discrete domains using spatial multi-omics data by combining the low-dimensional representation of biological profiles based on graph-based contrastive self-supervised learning. Our scalable method integrates multiple data modalities, such as RNA, protein, and H&E images, and predicts spatial domains within tissue samples. Through the integration of multiple modalities, Proust consistently demonstrates enhanced accuracy in detecting spatial domains, as evidenced across various benchmark datasets and technological platforms.

## 1 Introduction

Spatially-resolved multi-omics technologies enable the profiling of multiple omic measurements, such as the transcriptome and epigenome, in individual tissue sections, leading to an improved understanding of regulatory mechanisms along spatial coordinates [1–4]. These spatial technologies have already revolutionized our understanding of human tissue architecture and the impact on tissue architecture from disease [5–8]. Examples of these types of multi-omics technologies include measuring RNA and chromatin acc (Spatial ATAC–RNA-seq [3]) or measuring RNA and protein (DBIT-seq [1], or the 10x Genomics Visium Spatial Proteogenomics (SPG) [9] platform). In addition, the standard 10x Genomics Visium platform could also be viewed as a multi-omics technology through the integration of transcriptome data with a paired brightfield image after the tissues are stained with hematoxylin and eosin (H&E).

A standard step in the analysis of spatial multi-omics data is to identify discrete spatial domains. These domains can be further investigated for potential markers of tissue architecture corresponding to morphology [10] or unique niche-specific domains that might appear in complex diseases, such as cancer [11]. However, similar to single-cell data, it remains challenging to leverage supervised learning approaches to predict discrete spatial domains because of cell segmentation. Therefore, most existing tools used in practice today identify discrete spatial domains with either unsupervised or self-supervised learning approaches [12–15].

To identify discrete spatial domains from spatial multi-omics data, one approach is to ignore the spatial information entirely, consider only one of the omic data modalities (e.g. RNA), and apply unsupervised clustering methods used for single-cell, such as *k*-means [16], Louvain, and Leiden [17] algorithms. However, these approaches assume independence between the spatial coordinates and often lead to discontinuous or incoherent spatial domains [10]. A second approach is to continue with only one omic data modality, but to incorporate spatial information to account for the correlation of molecular information between the spatial coordinates. Some examples of these methods include (i) unsupervised learning approaches (BayesSpace [18], Giotto [19], STAGATE [20], CCST [21]) and (ii) self-supervised learning approaches (GraphST [15], SpaceFlow [22], ConGI [23], CAST [24]). In particular, the methods using contrastive self-supervised learning aim to maximize the similarity between adjacent spatial coordinates and dissimilarity between non-adjacent spatial coordinates, while also showing great promise in their ability to detect discrete spatial domains using only one data modality. A third set of tools aim to leverage more than one omic data modality, while also leveraging the spatial coordinates. SpaGCN [25] combines gene expression, spatial information, and histology image for spatial clustering using a graph convolutional neural network. However, this tool is designed to work only with hematoxylin and eosin (H&E) images. In this work, we aimed to address these limitations.

Inspired by the graph-based autoencoder and contrastive self-supervised learning frameworks for single omic data, here, we introduce Proust, a computationally scalable algorithm using contrastive self-supervised learning to predict discrete domains specifically designed spatial multi-omics data. We introduce an overview of the algorithm and compare the performance of our method to existing domain detection algorithms. By combining the lower-dimensional representation of multi-omic features that aggregate local tissue context through graph-based autoencoders, Proust identifies more biologically accurate and coherent tissue structures compared to existing state-of-the-art methods. In addition, we demonstrate how Proust can be used to detect discrete spatial domains in spatial tissue sections. Finally, we show how our method is computationally efficient and scalable in terms of memory and time, and we provide open-source software implemented in Python.

## 2 Results

### 2.1 Overview of Proust to detect spatial domains integrating multiple data modalities

Proust is a graph-based contrastive self-supervised learning framework to predict discrete spatial domains using spatial multi-omics data. For the purposes of clarity, we describe how Proust can detect spatial domains with two omic data modalities (RNA and proteins) (**Figure 1**). However, these ideas can be generalized to other types of multi-omics, for which we give examples of using RNA and brightfield images. Considering RNA and protein, Proust takes as input gene expression counts, multi-channel immunofluorescence (IF) images, and spot-level spatial position. The first step involves constructing a neighborhood graph structure based on the relative distance between spots. Next, graph-based convolutional autoencoders are trained separately for gene expression and extracted image features, aggregating genomic and protein information from neighboring locations. Furthermore, the framework uses contrastive self-supervised learning (**Figure S1**) to refine the latent embedding that maximizes similarities between adjacent spots while minimizing those between non-adjacent spots. The reconstructed gene expression and image features obtained from the graph-based decoder are used to extract the top principal components (PCs), which are used in conjunction with a model-based clustering algorithm, mclust [26], to identify spatial domains.

**Figure 1:**
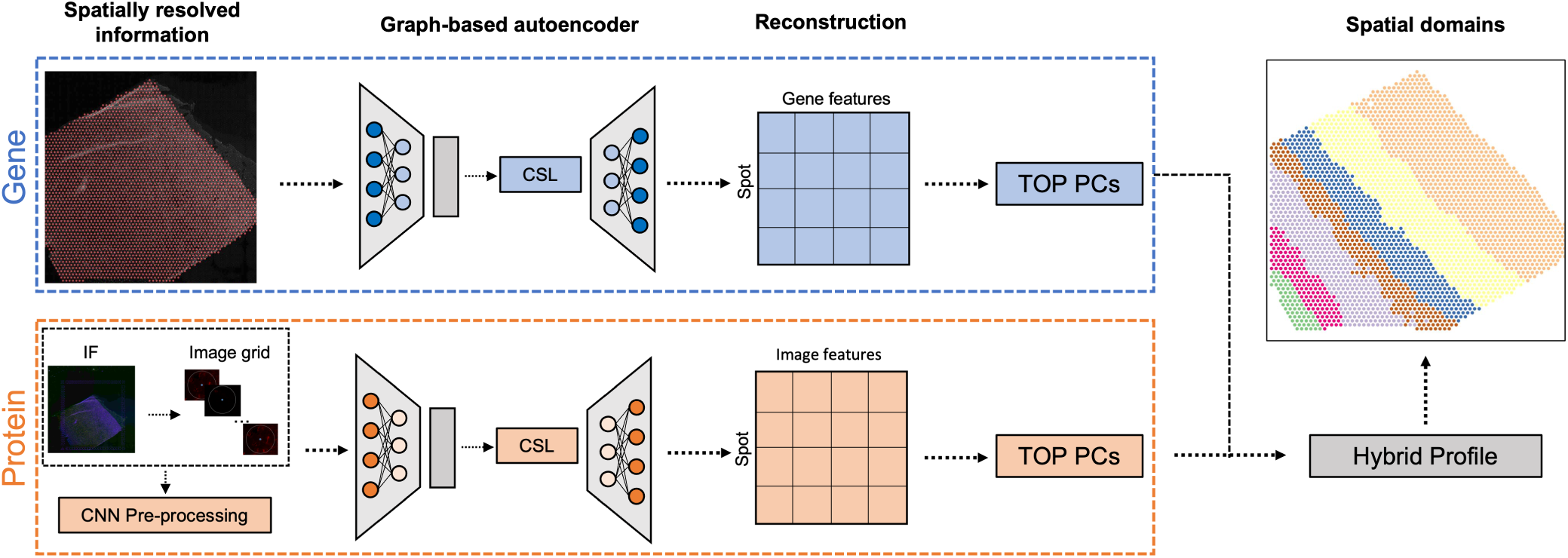
Overview of Proust for detecting discrete domains using spatial multi-omics data. For the purposes of clarity, we introduce Proust with two specific omic data modalities: RNA and protein, but these ideas can be generalized to other multi-omics, such as RNA and brightfield images. First, Proust constructs a graph structure based on the Euclidean distance between spatial coordinates. Next, graph-based convolutional autoencoders are trained separately for gene expression and protein information extracted from an immunofluorescence (IF) image. The latent embeddings are refined using contrastive self-supervised learning (CSL). The top principal components (PCs) from the reconstructed gene and image features are concatenated to create a hybrid profile for downstream clustering analysis.

### 2.2 Proust increases accuracy to detect spatial domains and marker genes

Proust was first applied to a CK-p25 mouse coronal brain tissue data set [27], measured on the 10x Genomics Visium Spatial Proteogenomics (SPG) platform (**Figure 2A**). This dataset includes transcriptome-wide gene expression and an IF image measuring *γ*H2AX, a protein involved in DNA repair. Proust’s latent embeddings showed distinct cortical domains that were well-separated from other brain regions (**Figure 2B**). The resulting predicted spatial domains from Proust were compared to other methods GraphST, SpaGCN, and STAGATE that employ graph convolutional neural networks. In one of the CK-p25 mouse brain tissue replicates, Proust identified spatial domain 4, 11, 9, and 5, corresponding to well-known mouse hippocampus (HPC) subfields, including CA1, CA2&3, dentate gyrus, and other HPC subfields, respectively, as per the Allen Brain Atlas [28] (**Figure S2**). On the other hand, the compared methods did not detect these finer granularity subfields the HPC, instead only identifying the HPC broadly (**Figure 2C**). As noted in Welch et al. [27], *γ*H2AX is associated with the enrichment of reactive microglia (RM). To assess the marker genes detected within the predicted spatial domains identified by Proust, subregional differential expression was assessed and a high expression of marker genes for RM were found including: *CST7*, *H2-D1*, *LGALS3BP*, and *LPL* (**Figure 2D**). This suggests that the integration of both the gene expression, with even just one protein of interest, can enable increased sensitivity to detect marker genes within domains with a finer spatial resolution.

**Figure 2:**
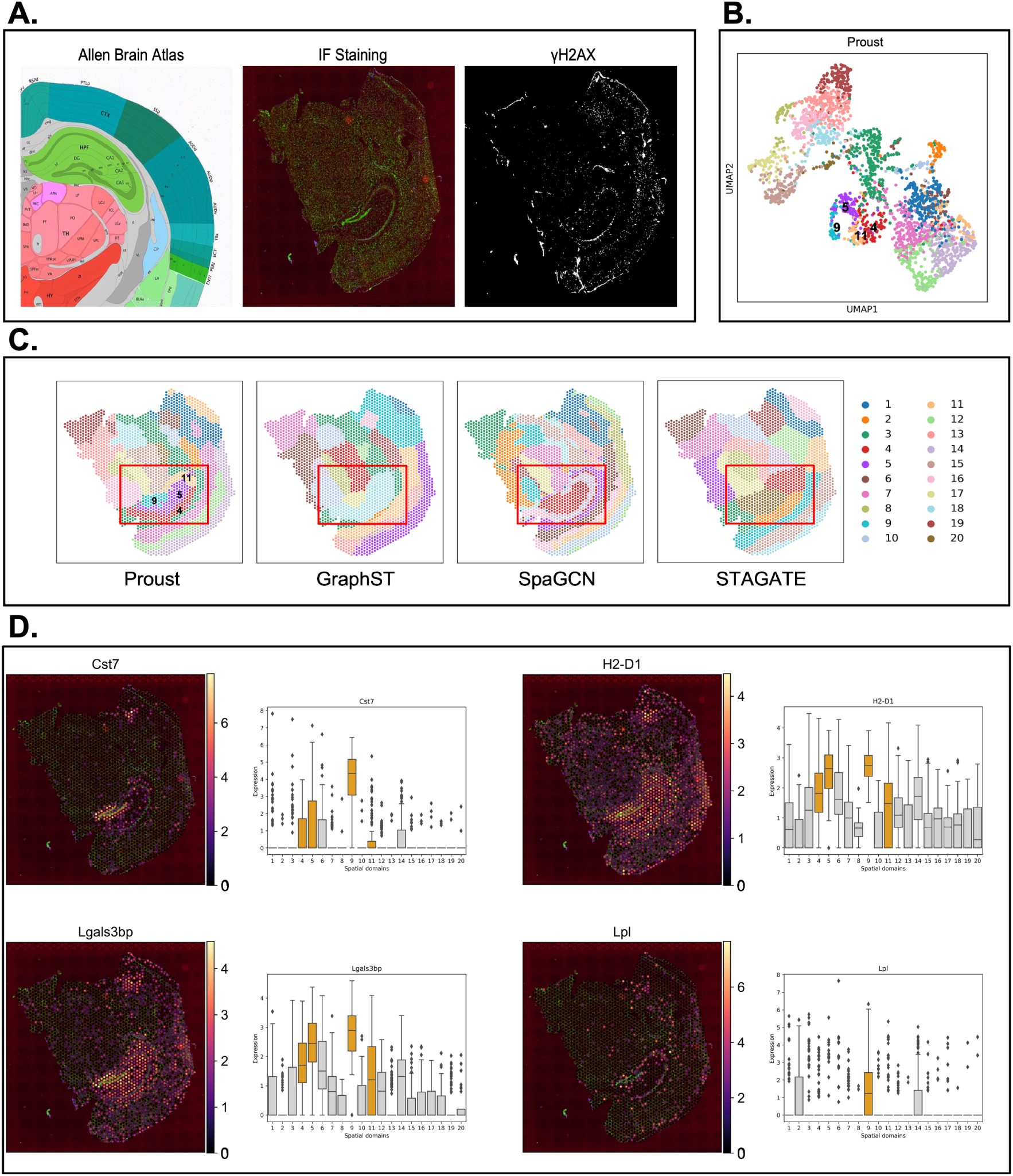
Proust improves the detection of hippocampal spatial domains in CK-p25 mouse coronal brain tissue by integrating gene expression with proteins of interest. **(A)** From left to right: (i) annotation of mouse hippocampus subfields from the Allen Reference Brain Atlas [28], (ii) merged DAPI and *γ*H2AX immunofluorescence images, and (iii) IF staining of *γ*H2AX. **(B)** UMAP representation of spots colored by spatial domains detected by mclust using Proust’s latent embeddings. **(C)** Predicted spatial domains by Proust, GraphST [15], SpaGCN [25], and STAGATE [20] with *k* = 20 domains. **(D)** Spatial expression level of four *γ*H2AX marker genes across the entire tissue slice and boxplots of corresponding marker genes stratified by *k*=20 domains identified by Proust. Hippocampal subregions are depicted in orange; other regions are depicted in gray.

Next, to evaluate the performance of the spatial domains detected by Proust, we used four Visium SPG human dorsolateral prefrontal cortex (DLPFC) brain tissue slices obtained from neurotypical donors [29], each with paired multiplexed IF images stained for nuclei and four cell types (**Figures 3A, S3**). The four tissue sections included manually annotated spatial domains for white matter (WM) along with six morphological domains (Layers 1-6 or L1-6). Using the adjusted Rand index (ARI) as a performance measure, we compared the similarity of these manual annotations to the predicted spatial domains. We evaluated the performance of Proust along with five existing clustering methods that are commonly used for spatial domain detection, namely GraphST [15], SpaGCN [25], and STAGATE [20], BayesSpace [18], and *k*-means [16].

**Figure 3:**
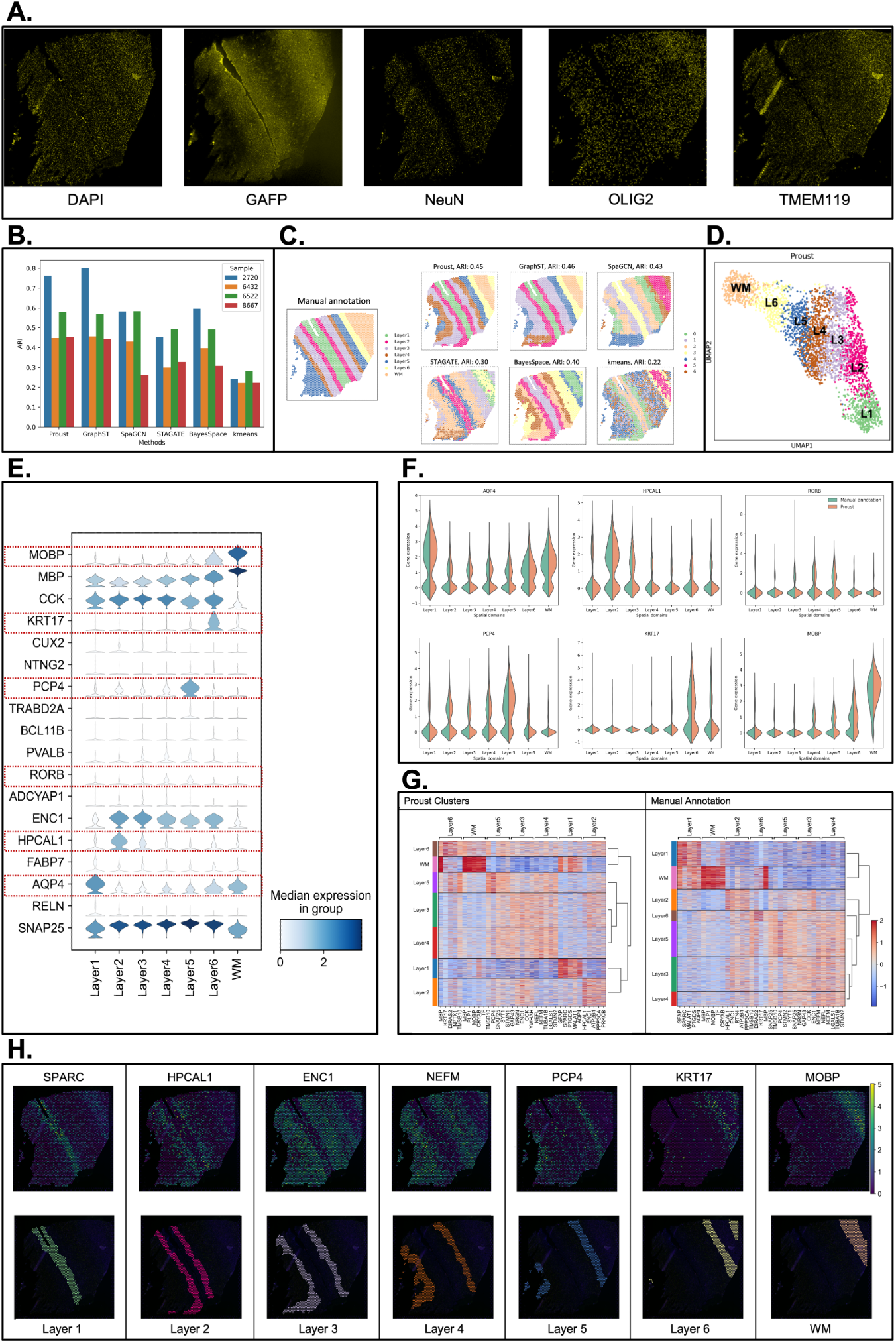
Proust improves the accuracy of predicting spatial domains compared to existing methods. Data from sample Br6432 from Visium SPG human DLPFC dataset [29], unless noted otherwise. **(A)** Immunofluorescence images of five protein channels: nuclei (DAPI), neurons (NeuN), oligodendrocytes (OLIG2), astrocytes (GFAP), and microglia (TMEM119). **(B)** Boxplot of adjusted Rand index (ARI) across four samples. **(C)** Manual annotation of tissue slice from donor Br6432 and predicted spatial domains by the six methods. Labels do not indicate corresponding biological layers assigned by the algorithms. **(D)** UMAP visualization of spots from donor Br6432 colored by Proust predictions. **(E)** Stacked violin plot of marker gene distribution for white matter and sub-layers of gray matter based on literature in each spatial domain assigned by Proust. Red rectangles highlighted in (F). **(F)** Violin plots of marker gene expression for Proust and manually annotated domains. **(G)** Heatmaps of the top 5 differentially expressed genes (centered and scaled) across layers from Proust and manual annotations. A dendrogram on the right shows hierarchical clustering. **(H)** Selected cluster-based marker genes expression and visualization of individual clusters identified by Proust. Layers were annotated according to the laminar organization indicated by the manual annotation.

It was found that the two GNN-based autoencoder and contrastive self-supervised learning frameworks (Proust and GraphST) resulted in the highest ARI, followed by SpaGCN, BayesSpace, STAGATE, with *k*-means showing the lowest ARI across the four tissue sections (**Figure 3B**). However, Proust, which integrated both RNA and protein data modalities, outperformed the other methods in recognizing more coherent and biologically meaningful gray and white matter layers (**Figures 3B-C, S4**).

Using Proust domains, we found that the UMAP (Uniform Manifold Approximation and Projection) [30], embeddings revealed separate cortical layers ordered in known morphological layers (**Figure 3D**). Using previously known marker genes for the cortical layers [10], Proust domains resulted in laminar-specificity with the known marker genes (*APQ4* for L1, *HPCAL1* for L2 and L3, *RORB* for L4, *PCP4* for L5, *KRT17* for L6, and *MOBP* for WM) (**Figures 3E, S5**), which also had a similar distributions of expression using marker genes using the manually annotated domains (**Figures 3F, S6**). However, using a data-driven approach, differentially expressed genes using Proust domains led to a more biologically meaningful hierarchical clustering of identified layers with white matter and L6 are grouped together, as opposed to the manual annotation, where instead white matter and L1 are grouped together (**Figure 3G-H**).

### 2.3 Proust flexibly weights data modalities to detect spatial domains

One advantage of Proust is the flexibility to weight multi-omics profiles (or data modalities), such as RNA and protein (**Figure 1**), to detect spatial domains particularly in the context of either healthy or disease tissue. Here, we use as an example a 10x Genomics Visium SPG dataset profiling human inferior temporal cortex tissue sections collected from individuals with late-stage Alzheimer’s disease (AD) [31]. In this dataset, the IF images contained five protein channels, namely nuclei (DAPI), amyloid-*beta* (Abeta), hyperphosphorylated tau (pTau), microtubule-associated protein 2 (MAP2), and astrocytes (GFAP) [31] (**Figures 4A, S7**). Also, the role of protein information is different from previous datasets, as some protein channels, are not useful to identify morphological domains. For example, Abeta is sparsely distributed throughout the tissue associated with AD pathology. Furthermore, Abeta and pTau are only detected at the protein level. In this section, we demonstrate how Proust can flexibly weight the mulitple data modalities to accurately identify spatial domains, even in diseased tissue.

**Figure 4:**
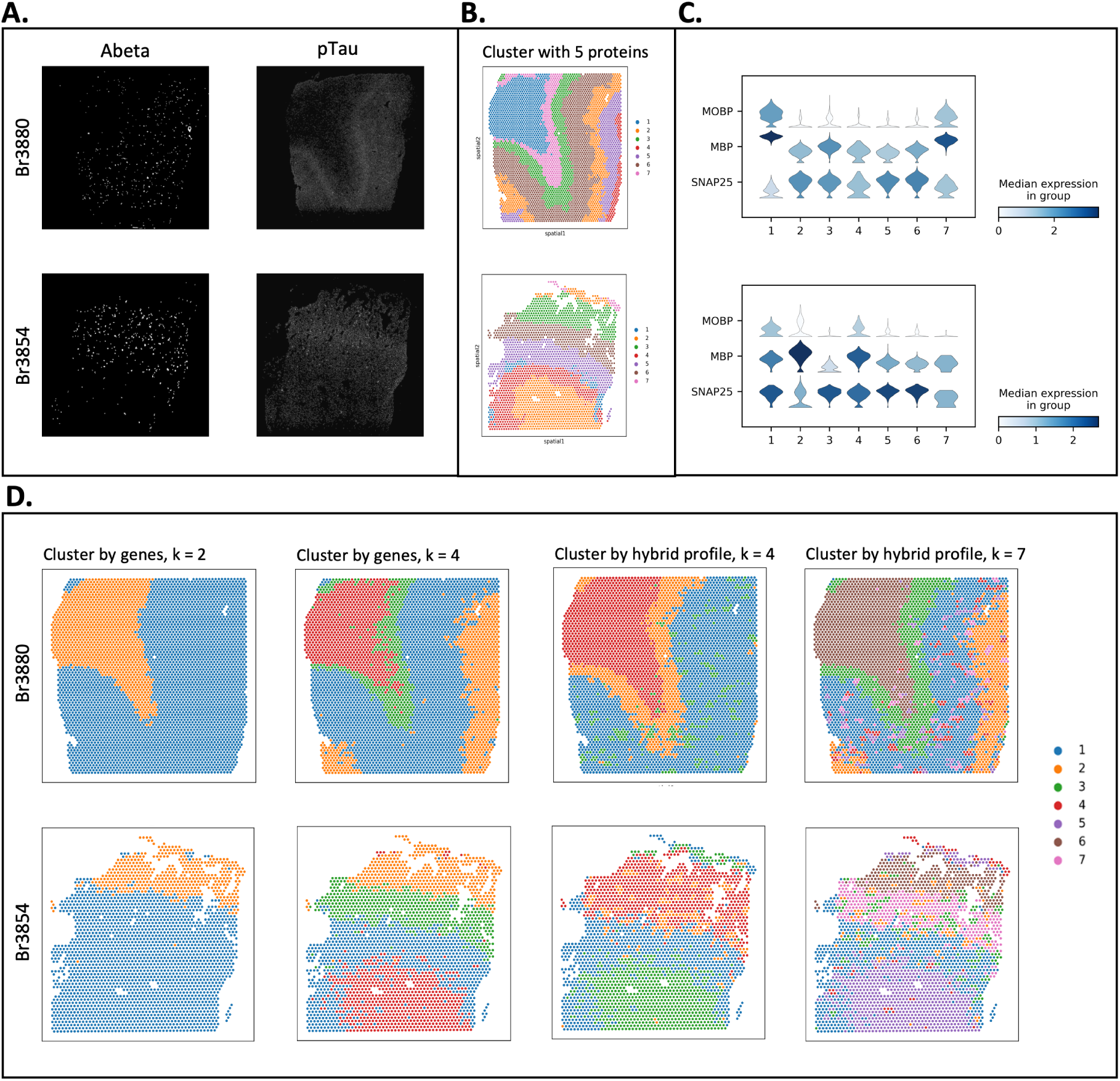
Proust achieves distinct spatial domain detection with different protein channels and weights assigned to transcriptomics and proteomics on Visium SPG human inferior temporal cortex tissue slices from donor Br3880 and Br3854. **(A)** Immunofluorescence staining images of Abeta and pTau. **(B)** Proust clustering result using five protein channels (DAPI, Abeta, pTau, MAP2, and GFAP), top 30 PCs from reconstructed gene expression, top 5 PCs from reconstructed extracted image features, and *k* = 7 clusters. **(C)** Stacked violin plot of distribution of marker genes (*MOBP* for oligodendrocytes/WM, *SNAP25* for neurons/gray matter) in each spatial domain assigned by Proust. **(D)** Proust clustering result using two protein channels (Abeta and pTau). The first two columns show clustering results using transcriptomics only when *k* = 2 and *k* = 4 clusters, respectively. The last two columns show clustering results using a hybrid profile of transcriptomics and proteomics, with the top 10 PCs from reconstructed gene expression and the top 10 PCs from reconstructed extracted image features when *k* = 4 and *k* = 7 clusters, respectively.

For example, Proust can use all five protein channels to create a hybrid profile (top 30 PCs from the reconstructed gene expression and the top 5 PCs from the reconstructed extracted image features) to identify spatial domains (**Figures 4B, S8**), which can be used to visualize marker genes associated with the white and gray matter (**Figure 4C**). Alternatively, we can compare predicted spatial domains from Proust (i) only using gene expression (and ignoring proteins) (**Figure 4D**, columns 1 and 2 using *k*=2 or *k*=4, respectively) and (ii) using gene expression and only two protein channels (Abeta and pTau) (**Figure 4D**, columns 3 and 4 using *k*=4 or *k*=7, respectively).

In the later case, Proust can weight the RNA and protein information separately by controlling the number of PCs extracted from each data modality. This can lead to detected spatial domains corresponding to known morphology in healthy tissue, pathologies associated with disease, or both. For instance, in the tissue slice from Br3880, Proust identified spotty areas (cluster 3) and a layer (cluster 2), which are visually correlated to Abeta-and pTau-captured areas when clustering by a hybrid profile for *k* = 4 clusters. In contrast, spatial domains corresponding to only cortical layers were detected when clustering by genes alone with the same number of clusters (**Figures 4D, S9**). Upon increasing the number of clusters to 7, Proust distinguished additional sub-layers within the gray matter with higher precision while retaining regions associated with Abeta and pTau. These results demonstrated that Proust is able to detect relevant spatial domains of interest by leveraging different protein channels and flexibly adjusting the number of PCs from each data modality into the hybrid profile. Finally, it was also observed that Proust’s performance improved when the broad and connected spatial pattern is evident in IF images to complement expression information.

### 2.4 Proust accurately detects spatial domains with expression and histology images

Next, we demonstrate how Proust can be generalized to other types of multi-omics, specifically with gene expression and H&E brightfield images, rather than IF staining, measured on the 10x Genomics Visium Spatial Expression platform [32]. To evaluate the performance of Proust, we compared the predicted spatial domains to results from five existing clustering algorithms on *N* =12 Visium human DLPFC tissue slices that have manually annotated spatial domains to be used as a gold standard [10]. The three RGB channels were included separately at the pixel level in the image feature extraction steps and autoencoder model training. Proust achieved the highest ARI with a median (across *N* =12 tissue sections) value of 0.56 and exhibited comparable performance to that of GraphST (median ARI = 0.53) (**Figure 5A**). The predicted spatial domains for 11 out of 12 samples obtained ARIs higher than 0.50, suggesting that Proust was effective when applied to histology images. Also, the cortical layers segmented by Proust were more biologically consistent with manual annotations including having greater spatial contiguity than the other methods, which tended to be more disconnected (**Figures 5B, S10**). In particular, Proust was able to identify thinner layers, such as Layer 2, and provided coherent sub-layers within the gray matter, as demonstrated in samples 151509 and 151674.

**Figure 5:**
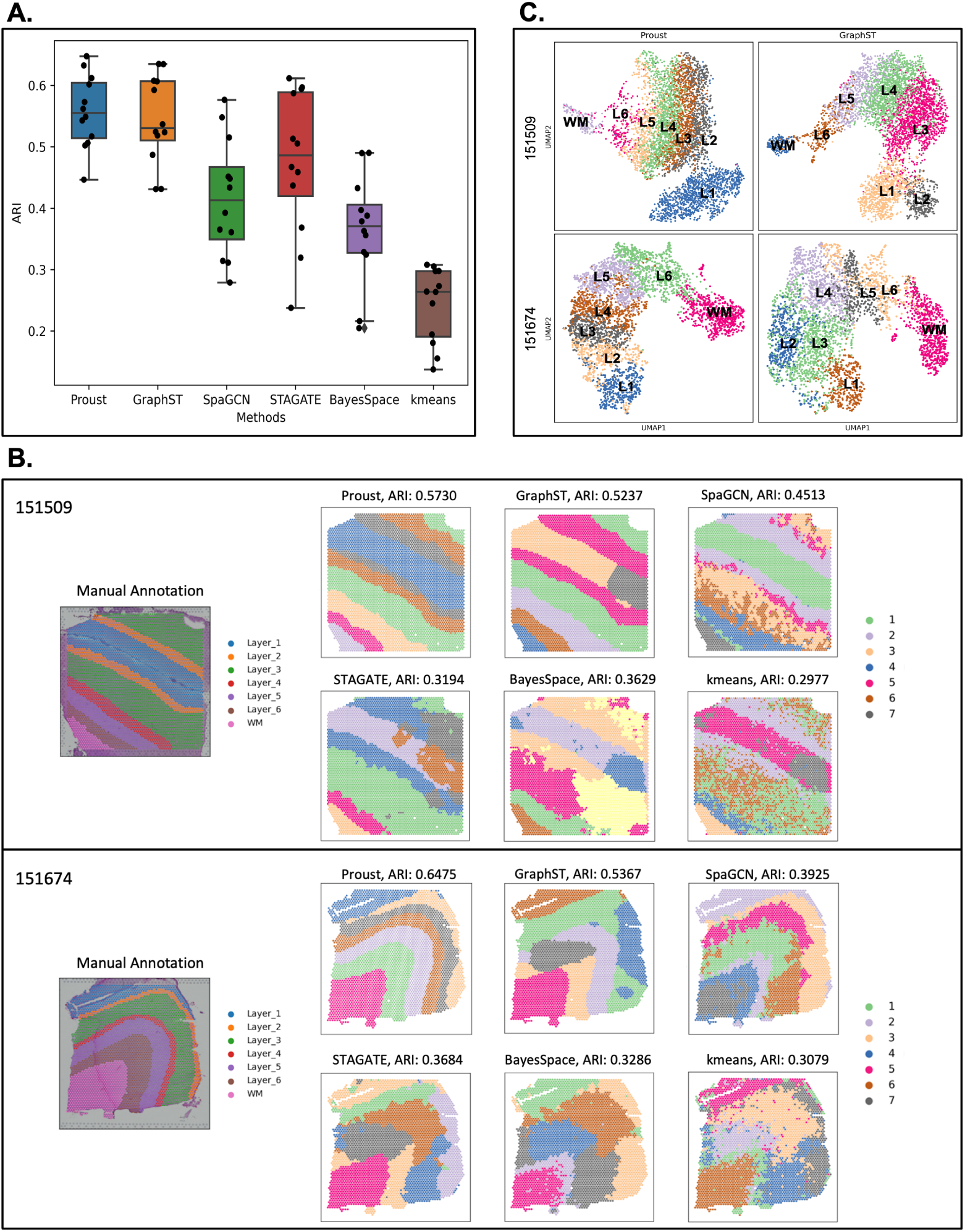
Evaluating and comparing the performance of Proust in layer segmentation with other popular existing methods on the Visium human DLPFC dataset that contains H&E images. **(A)** Boxplot of clustering accuracy in 12 DLPFC samples across Proust and five other existing methods based on adjusted rand index (ARI). **(B)** Manual annotation of tissue slices 151509 and 151674 and spatial domains assigned by the six methods. **(C)** UMAP visualization of reduced dimensions from Proust and GraphST for 151509 and 151674.

Although Proust and GraphST yielded similar ARIs, the UMAP plots of latent embeddings from Proust demonstrated more meaningful functional similarity between adjacent clusters (**Figure 5C**). These results suggest that Proust can also effectively extract histology image features that distinguish neighboring layers and refine the detection of coherent spatial regions and functional domains.

## 3 Discussion

Proust is a novel framework that utilizes spatially-resolved transcriptomics, immunofluorescence staining images for protein channels, and spatial location information for the identification of spatial domains in 2-D tissue slices. Proust effectively reconstructs biological features that aggregate spot-level neighboring information through GCN-based autoencoders and contrastive self-supervised learning. By combining the significant spatial variance explained by transcriptomics and proteomics, Proust can capture complex dependencies and spatial patterns, allowing for spatial segmentation of tissue structures with high accuracy. Proust outperforms five popular existing methods in layer segmentation on human and mouse brain datasets generated from the 10x Visium platform, identifying biologically meaningful layers consistent with manual annotations in human dorsolateral prefrontal cortex regions and improving the detection of mouse hippocampus structures enriched with proteins of interest. The newly developed STACI method, which jointly analyzes spatial transcriptomics and chromatin imaging data with over-parameterized graph-based autoencoders, is not evaluated here as it is not available as a package at the time of conducting and writing this project and has been shown to fail to identify HPC sub-structures in a Visium mouse brain coronal section dataset [as per 33].

Additionally, Proust is a highly adaptable framework that allows users to adjust the weights assigned to gene expression and protein information according to their needs. By selecting different protein channels and adjusting top PCs of different data modalities used in the downstream clustering step, Proust can better capture the spatial domains enriched with specific proteins of interest. For instance, when assigning higher weights to IF protein images, Proust can detect regions that contain sufficient levels of protein associated with disease within identified broad biological layers, as shown in the Visium SPG human inferior temporal cortex dataset. However, we acknowledge the need to develop a more automatic weight adjustment procedure in Proust. Furthermore, Proust can incorporate histology information in addition to spatial transcriptomics when IF images are unavailable in the dataset, thus making it a versatile tool for spatial analysis using various image types. Overall, Proust’s flexibility allows users to fine-tune their analysis and obtain more accurate results tailored to their specific research questions and data types.

The use of Proust in the current project has some limitations and caveats. One assumption is that adjacent spots or tissue areas have similar biological profiles for spatially resolved gene expression and protein information. This assumption of the universal law of geography led to the implementation of graph-based neural networks for feature learning for both data modalities. However, for some proteins, such as Abeta, expression may be disconnected on the spot level, especially in late stages of progressive diseases, potentially making it challenging for Proust to accurately identify all regions where the protein occurs, as individual spots may absorb dissimilar neighboring information during model training. To address this issue, a potential solution is to enhance the graph structure with weighted edges based on the similarity of spot-level protein information or to incorporate inter-modality contrastive learning to maximize the mutual information between gene expression and proteomics [23, 34]. We are also interested in exploring the use of statistical inference here as a way to explore the uncertainty of the predicted spatial domains. Additionally, since Proust is designed for spot-level information extraction when spatial coordinates for spot centroids are provided in 10x Visium datasets, it fails to recover pixel-level protein information from IF images unless strong signals of associated spatially resolved gene expression are present. We would like to expand and test this current method to datasets from other platforms that may include subcellular spatially-resolved transcriptomic and proteomic data. We also observed that image noise reduction is a crucial preprocessing step before utilizing Proust, as the method may be affected by pixel values that result from technical artifacts. The average computational time for analyzing the four tested datasets is approximately 8-12 minutes per tissue sample on MacOS 13.0, using an Apple M1 Pro chip and 32GB memory. We anticipate that datasets with more spots and protein channels will require more memory and processing time. Thus, we recommend enabling the GPU option during model training to achieve quicker performance.

## 4 Methods

### 4.1 Data availability

#### 4.1.1 CK-p25 mouse coronal brain data set

The CK-p25 mouse coronal brain tissue data set [27] was measured on the 10x Genomics Visium SPG platform [35]. In this dataset, the IF images captured *γ*H2AX, a protein involved in DNA repair. To enhance the *γ*H2AX gray-scale image and reduce noise, we established a threshold of 6 standard deviations above the mean pixel values and applied a max filter with a 10 *×* 10 image box to facilitate the detection of protein-rich regions by Proust.

#### 4.1.2 DLPFC Visium SPG

The human DLFPC brain tissue was measured on the 10x Genomics Visium SPG platform [29].

#### 4.1.3 ITC

The human inferior temporal cortex tissue sections collected from individuals with late-stage Alzheimer’s disease (AD) were profiled with the 10x Genomics Visium SPG platform [31]. In this dataset, the IF images contained five protein channels, namely nuclei (DAPI), amyloid-*beta* (Abeta), hyperphosphorylated tau (pTau), microtubule-associated protein 2 (MAP2), and astrocytes (GFAP). We used the preprocessed grayscale images from VistoSeg [36] and enhanced them in a manner similar to that employed in analyzing the Visium SPG mouse CK-p25 brain tissue dataset. To amplify the influence of protein information on spatial segmentation, we decreased the number of top PCs for transcriptomics to 10 while increasing those for proteomics to 10 from the default setting to create a hybrid biological profile.

#### 4.1.4 DLPFC Visium H&E

The human DLFPC brain tissue was measured on the 10x Genomics Visium Spatial platform with H&E images [10].

### 4.2 Spatially-resolved gene expression preprocessing

We evaluated the performance of Proust by testing it on four Visium datasets generated from the 10x Genomics platform [35]. In Visium, the capture area measures 6.5 * 6.5 mm. The spots have a diameter of 55 µm (equivalent to an area of 2,375 µm^2^) and are spaced 100 µm center-to-center in a honeycomb pattern (i.e., each spot is surrounded by six adjacent neighbors). Each tissue slide contains a total of 4992 sequenced spots, capturing around 35,000 genes. The raw gene count matrix is generated by aligning Visium sequencing data with the fiducial frame on the histology or immunofluorescence (IF) staining image of the tissue slice. Two-dimensional (2D) spatial positions are provided as pixel coordinates of spot centroids. The first three test datasets were generated using Visium SPG, which include multiplexed immunofluorescence images of proteins of interest from the corresponding tissue area. Additionally, we evaluated the transferability of Proust on a Visium dataset containing hematoxylin and eosin (H&E) images of the tissue slice.

To prepare the datasets for analysis, spots outside the tissue area are removed before applying Proust’s standard preprocessing steps for gene expression using the SCANPY package [37]. The spatially resolved gene expression information is represented by an *N × G* matrix for *N* spots and *G* types of genes, with spot centroid locations indicated by 2-D spatial coordinates (*x, y*). Spike-in and mitochondrial genes are filtered out, and those expressed in fewer than three spots are excluded. Raw gene counts are then (1) normalized by library size, (2) log-transformed, and (3) scaled to unit variance, with any values exceeding a standard deviation of 10 clipped. The top 3,000 highly variable genes (HVGs) are selected as input for the subsequent model training.

### 4.3 Image feature extraction

Protein density can vary across different spatial regions, offering valuable information that can be integrated with spatial transcriptomics to enhance tissue architecture inference. Spot-level protein information is obtained by first splitting the full-resolution image into *d × d* circumscribed grids centered on each spot centroid, where *d* is the ceiling rounding for the spot diameter. To improve computational efficiency, we reduce the size of each image grid to 48*×*48 pixels for 10x Genomics datasets, as the spot diameter is usually larger than or around 100 pixels. This downsizing is achieved by using the OpenCV Python package [38], where the updated pixel value of the new interpolated image is calculated as a weighted average of input pixel values from a sliding window grid in the original image. The grid size is determined by the ratios of the original image’s width and height to those of the resized image. To ensure that different modalities of data can be jointly analyzed and achieve optimal model performance in identifying spatial patterns using gene expression and protein data, pixel values are normalized to the same range as the preprocessed gene expression prior to model training.

Next, we implement a simple convolutional neural network (CNN) autoencoder to extract protein features from interpolated image grids. The model takes the resized image grid *I* for each spot as input. The encoder consists of two fully-connected inner layers with a kernel size of 5 *×* 5, each followed by an average pooling layer with a kernel size of 2 and stride of 2. The decoder consists of two transposed convolutional layers to reconstruct latent representations and images with the same dimensions as those of the intermediate output from the encoder. A ReLU (Rectified Linear Unit) activation function is applied after each convolution step, which allows the CNN model to learn complex and abstract features from images by introducing non-linearity to the output. We train this model by minimizing the self-reconstruction loss of image grids using the following equation:

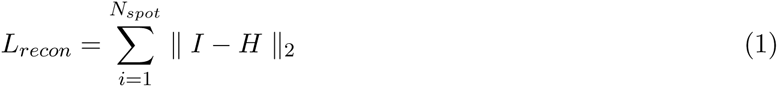

where *H* is the reconstructed image grid. We use the Adam optimizer to minimize the reconstruction loss with an initial learning rate of 1e-3. The default number of iterations is set to 800. The latent embeddings produced by this CNN-based autoencoder capture continuous spot-level spatial patterns for each protein channel instead of describing pixel-level intensity. This approach enables us to focus on the broad changes in protein distribution within the stained tissue slice, which complements our analysis of spatially-resolved gene expression.

### 4.4 Graph convolutional autoencoder

#### Graph structure based on spatial information

Incorporating spatial information along with biological features is crucial for identifying coherent spatial domains, as it allows for considering neighboring information from the nearest spots. To leverage the spatial information, Proust first converts the spot-level spatial coordinates into an undirected neighbor graph *G* = (*V, E*) with a pre-defined (or user-defined) neighbor number *k*. Here, *G* represents the graph, *V* represents spots, and *E* represents connected edges between each pair of spots (*i, j*) *∈ V*. The structure of the graph *G* based on spatial proximity is measured using the adjacency matrix *A ∈* ℝ*^N×N^*, where *N* is the number of spots. For a given spot *i*, the adjacency matrix assigns a value of 1 to *A_ij_*if spot *j* is among the *k*-nearest neighbors selected based on the Euclidean distance; otherwise, it assigns a value of 0. To normalize the influences across spots, a symmetric normalized Laplacian matrix 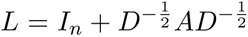 is constructed, which also takes into account the information from each spot itself. The degree matrix *D* is a diagonal matrix with elements 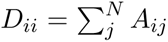 being the number of edges (i.e., neighbors) attached to each spot. When constructing the graph structure, we tested on *k* = 1, 2, 3, 4, 5, or 6, since there are six contiguous neighbors surrounding each spot *i* in 10x Genomics datasets. Proust yields the best performance in spatial clustering when *k* is set to 3 for most tested datasets.

#### GCN-based autoencoder to reconstruct transcriptomic and proteomic features

We construct an autoencoder using a graph convolutional network (GCN) to learn a latent representation *Z_i_* for spot *i* by aggregating information from neighboring spots *j*’s that share similar biological profiles. In this approach, the GCN-based autoencoder takes preprocessed biological features *X* and spatial information stored in *L* as input and outputs the reconstructed spot-level feature matrix *H*.

The (*t* + 1)-th layer representation in the encoder for spot *i* is constructed using a trainable weight matrix 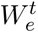 and a trainable bias vector 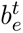, with a non-linear activation function ReLU denoted by *σ*(*·*). The encoder representation is formulated as follows:

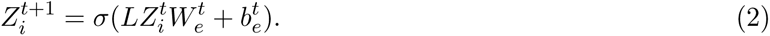

We denote *Z*^0^ as the original feature matrix *X* as input and *Z* as the final output of the encoder. The latent representation *Z* is then fed into a decoder to reconstruct the feature space *H* iteratively. The (*t −* 1)-th layer representation in the decoder is constructed using a trainable weight matrix 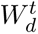 and a trainable bias vector 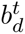, with the same activation function ReLU. The decoder representation is formulated as follows:

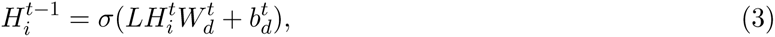

where the first inner layer 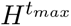 in the decoder is set as the final latent space *Z* from the encoder, and *H*^0^ is the reconstructed feature matrix *H*.

The autoencoder for gene expression data takes the preprocessed gene counts as input. On the other hand, the autoencoder for protein information takes the extracted image features as input, which are flattened for each channel before being fed into the encoder. This image autoencoder is three-dimensional, where the first dimension is the number of observations (i.e., spots), the second dimension is the number of protein or image (i.e., RGB channels for H&E histology images) channels, and the third dimension is the protein features. By training separate autoencoders for transcriptomic and proteomic data, the model can capture distinct patterns and features specific to each modality, which can then be integrated before the downstream clustering procedure.

### 4.5 Contrastive self-supervised learning

Proust addresses the challenge of distinguishing spots from similar spatial domains by implementing contrastive self-supervised learning (CSL) adapted from Deep Graph Infomax [39] to learn more discriminative reconstructed biological features (Figure S1). Contrastive self-supervised learning is a method for training deep neural networks to extract representations of data by comparing similar and dissimilar pairs of samples without relying on labeled data. This refinement step enables the learning of attributes that are common between data groups and attributes that separate one data group from another. In the data augmentation part, we generate a corrupted neighboring graph structure (*X^′^, A*), as opposed to the original structure (*X, A*), by randomly shuffling biological features while preserving the distance-based graph representing the spatial proximity between pairs of spots. We then feed the real and corrupted graph structures into a shared GCN-based autoencoder to obtain corresponding spot-level latent representations *Z* and *Z^′^*, respectively. A local neighborhood context *S_i_* for a given spot *i* is summarized with a read-out function defined as follows:

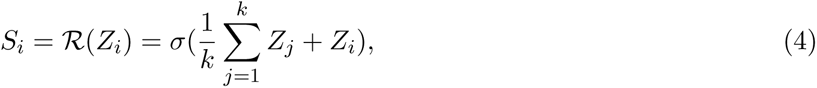

where *k* is the number of nearest neighbors for each individual spot. To maximize the mutual information between spot embeddings and local neighborhood embeddings, we calculate a discriminative score for context-spot representations pairs using a simple bilinear scoring function as the following:

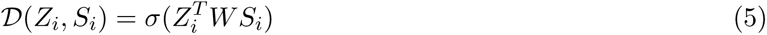

where *W* is a trainable scoring matrix and *σ*(*·*) is the logistic sigmoid function. A positive pair formed by *Z_i_* with the corresponding real local representation *S_i_* (or corrupted 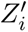 with 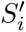) will be assigned a higher probability score, whereas a negative pair formed by *Z_i_* with the corrupted local representation 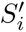 (or corrupted 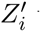 with real local representation *S_i_*) will be assigned a lower probability score. CSL refines the final latent embeddings from the encoder before reconstructed layers in the decoder. We train the GCN-based autoencoder with CSL by minimizing an overall loss combined from the self-reconstruction loss of the autoencoder and contrastive loss:

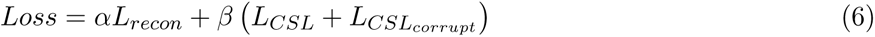

where

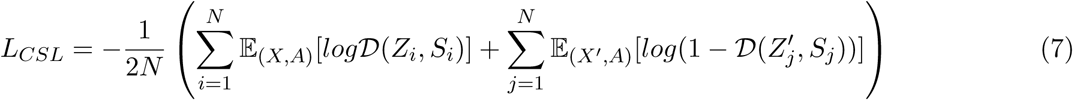

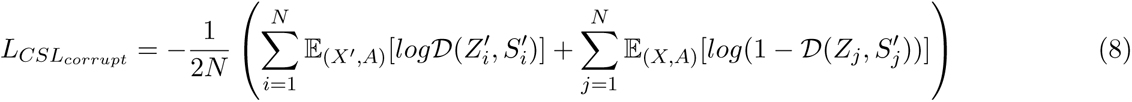

Hence, this supplementary process helps the GCN-based autoencoder to reconstruct the updated biological features, which bring similar spots together and differentiate dissimilar ones while preserving the fundamental information from the original input matrix. The Adam optimizer is utilized for training this deep learning architecture with an initial learning rate of 1e-3. The default number of iterations is set to 600.

### 4.6 Clustering and refinement

After training GCN-based autoencoders with preprocessed transcriptomic and proteomic features, Proust extracts the top principal components (PCs) from the reconstructed gene expression and extracted protein features separately. These components are then concatenated to create a hybrid biological profile for cluster assignment using a non-spatial clustering algorithm called mclust [26]. The default number of PCs used for transcriptomics is 30, and the default number of PCs used for proteomics is 5. Users can adjust the number of PCs to give adjusted weights for the two data modalities based on their needs. We pre-set the cluster count in mclust to correspond with the number of clusters in datasets that have manual annotations. For datasets without prior knowledge of the correct cluster count, we evaluated various cluster numbers and chose the one yielding the highest Silhouette score [40]. After clustering, Proust also offers an optional refinement step in which a given spot *i* is relabeled to the most common spatial domain of its *r* nearest surrounding spots. The default setting for *r* is 10.

### 4.7 Evaluation and comparison

#### 4.7.1 Adjusted Rand Index

To evaluate the performance of the Proust framework, we use the Adjusted Rand Index (ARI) to measure the agreement between identified spatial domains and manual annotation for individual tissue slices. Let *Ŷ* represent the assigned spatial clusters and *Y* represent the ground truth of clusters of *N* spots. Then,

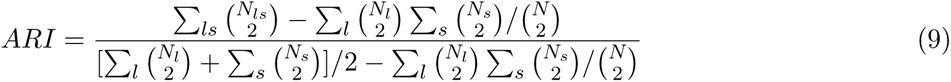

where *l* and *s ∈ m* clusters, 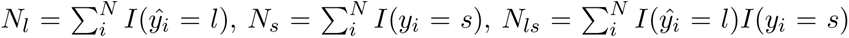. *I*(·) is an indicator function that follows *I*(*a* = *b*) = 1 when *a* = *b*, otherwise 0. The ARI ranges from 0 to 1, wherein a higher value indicates a better match between clustering results with the manual annotation.

#### 4.7.2 Existing methods

We benchmark Proust against the following existing methods for spatial domain detection:

##### GraphST

GraphST is a graph contrastive self-supervised learning framework that incorporates spatial information and gene expression for spatial clustering [15]. We followed the tutorial with default parameter settings and set r = 50 during refinement.

##### SpaGCN

SpaGCN combines gene expression, spatial information, and histology image for spatial clustering using graph convolutional neural network [25]. We followed the tutorial to use SpaGCN with the default parameter settings. The ‘histology’ option is disabled when no histology information is available in the dataset.

##### STAGATE

STAGATE is a method that utilizes an autoencoder and graph attention mechanism to learn a latent representation by incorporating spatial information and gene expression [20]. We followed the tutorial with default parameter settings and set the cell type-aware module ‘alpha’ to 0.

##### BayesSpace

BayesSpace is a spatial clustering method that utilizes a Markov random field with a Bayesian framework. The method assigns greater importance to adjacent spots by incorporating a prior that considers the spatial proximity of the spots [18]. We followed the tutorial with default parameter settings and set the number of iterations ‘nrep’ to 10000.

##### *k*-means

*k*-means is a clustering algorithm that partitions a set of *n* data points into *k* clusters, where each data point belongs to the cluster with the nearest centroid profile [16]. As the only method compared that is not specifically designed for ST data, *k*-means is used as the baseline comparison. We followed the tutorial from the bluster R/Bioconductor package with default parameter settings [41].

## 5 Back Matter

### 5.1 Code Availability

Proust is freely available on GitHub at github.com/JianingYao/proust. Code to reproduce all preprocessing, analyses, and figures in this manuscript is available from github.com/JianingYao/proust_paper. We used Proust version 1.0 for the analyses in this manuscript.

## 5.2 Acknowledgements

We would like to acknowledge Erik Nelson, Sang Ho Kwon, and Kasper Hansen for their helpful comments, feedback and suggestions on the methodology and manuscript. We would like to acknowledge Kristen Maynard for the manual annotations of the Visium H&E images. Erik Nelson, Sang Ho Kwon, and Kristen Maynard are from Johns Hopkins School of Medicine and Lieber Institute for Brain Development. Kasper Hansen is from the Johns Hopkins Bloomberg School of Public Health, Department of Biostatistics. We also thank the maintainers of the Joint High Performance Computing Exchange (JHPCE) compute cluster at Johns Hopkins Bloomberg School of Public Health for providing essential computing resources.

## 5.6 Abbreviations

Abeta: amyloid-*beta*
AD: Alzheimer’s disease
ARI: adjusted Rand index
DLPFC: dorsolateral prefrontal cortex
H&E: hematoxylin and eosin
IF: immunofluorescence
PCA: Principal component analysis
pTau: hyperphosphorylated tau
SRT: spatially-resolved transcriptomics
Visium SPG: 10x Genomics Visium Spatial Proteogenomics platform
UMAP: uniform manifold approximation and projection
WM: white matter

## 5.7 Author Contributions

JYao developed the Proust framework, performed analyses, created figures, and drafted text. JYu implemented Proust in a software package and provided input on the methodological framework, and text. BC provided advice on the architecture Proust, and edited the text. SCP and KM provided input on the biological interpretation of the data used, figures, and text. SCH supervised the project; provided input on the methodological framework, software implementation, analyses, figures, and text; and drafted text. All authors approved the final manuscript.

## Supplementary Figures

**Figure S1:**
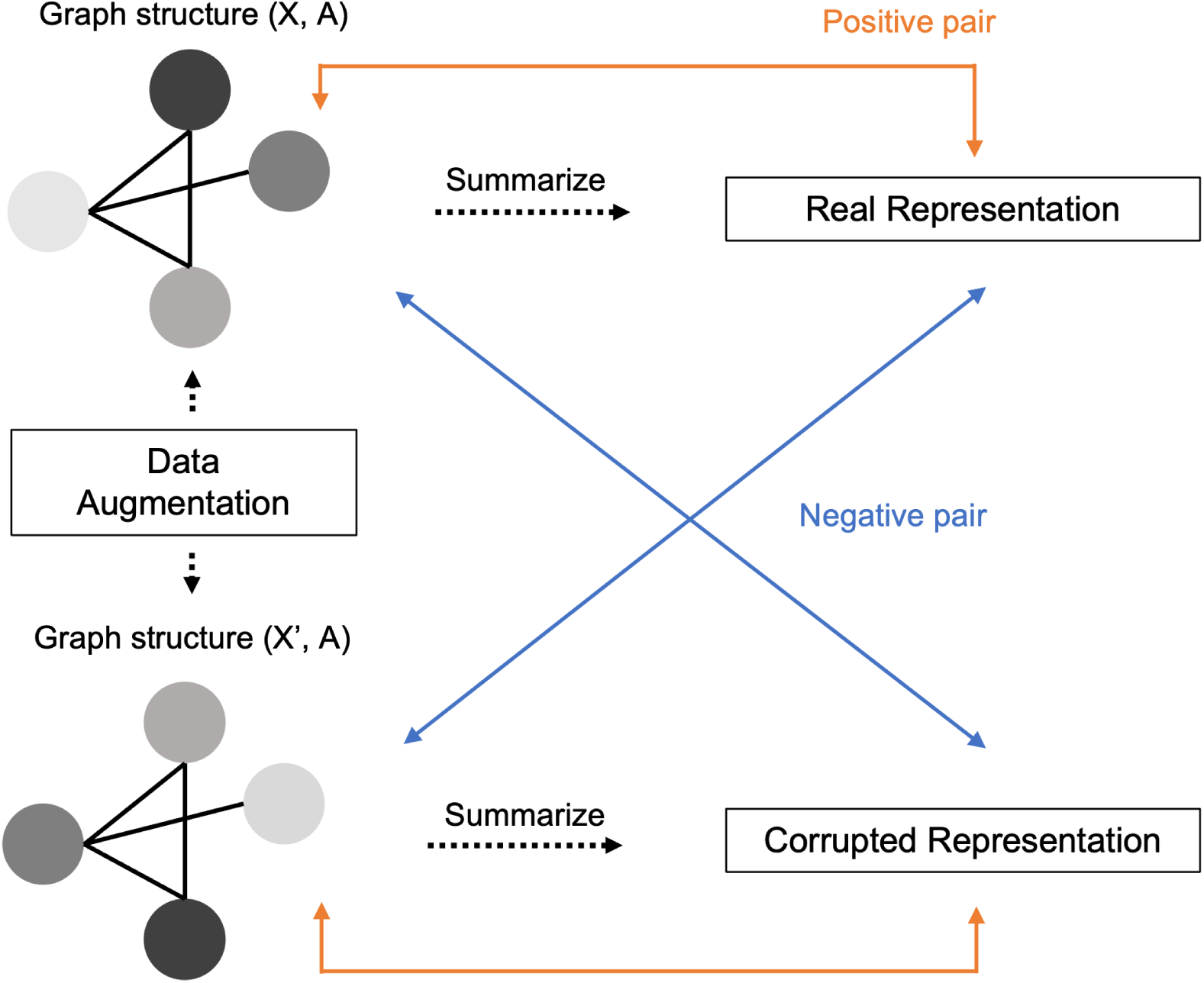
Contrastive self-supervised learning. Contrastive self-supervised learning is illustrated in this figure, demonstrating the refinement of latent representations during training of a graph-based autoencoder model. In the data augmentation step, biological features are randomly shuffled while preserving the distance-based graphs connecting each observation. Real and corrupted local representations are then summarized from these two sets of graph structures using a read-out function. A discriminative score for each pair of spot-patch representations is calculated during each iteration, comparing the spot-level latent embeddings with the summarized local context, respectively.

**Figure S2:**
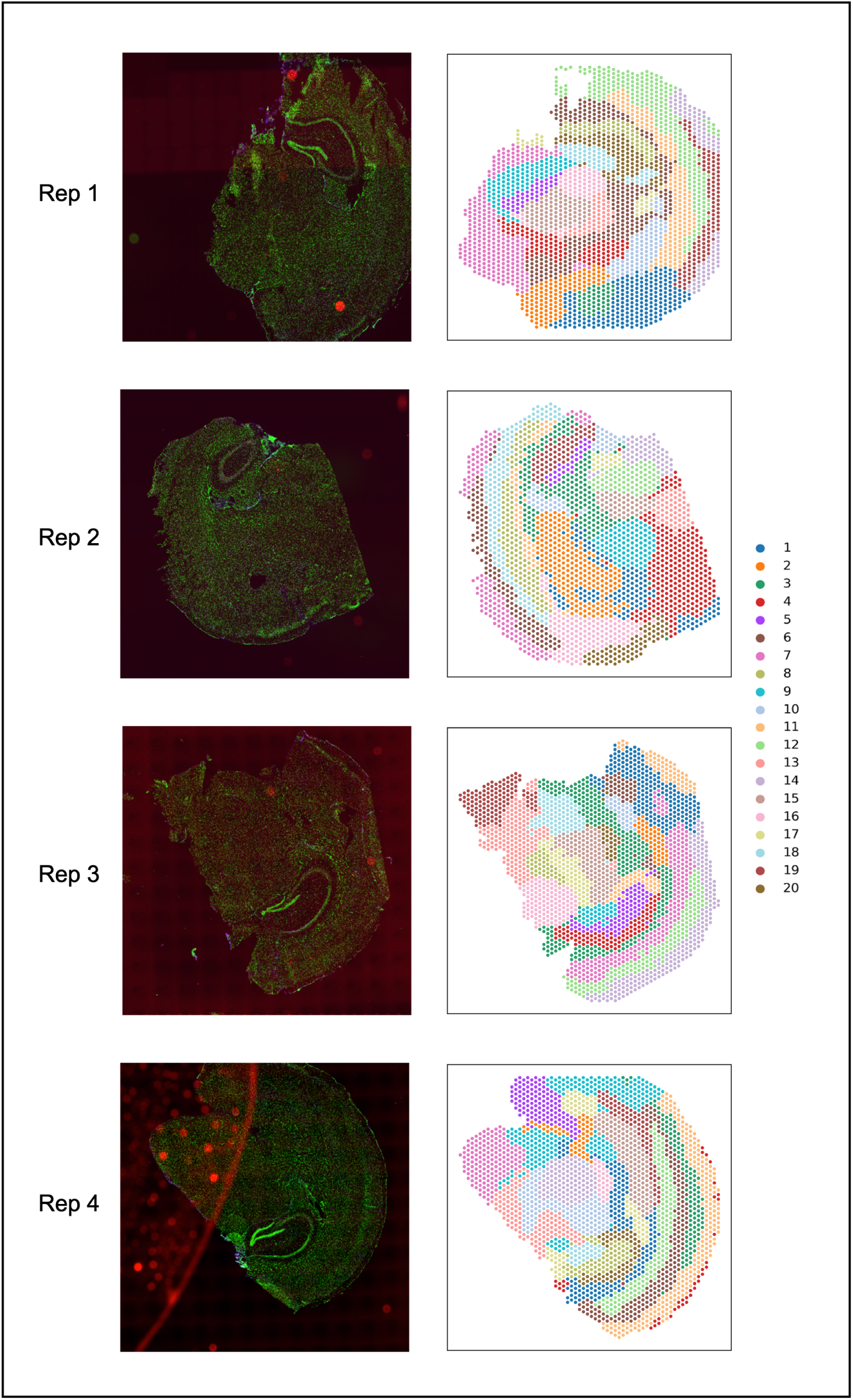
CK-p25 mouse coronal brain tissues measured on the Visium SPG platform across four tissue replicates. For each of the four tissue replicates (rows), the IF staining images of *γ*H2AX protein reproduced from Welch et al. [27] (left column) and the spatial domains detected by Proust for *k*=20 domains (right column).

**Figure S3:**
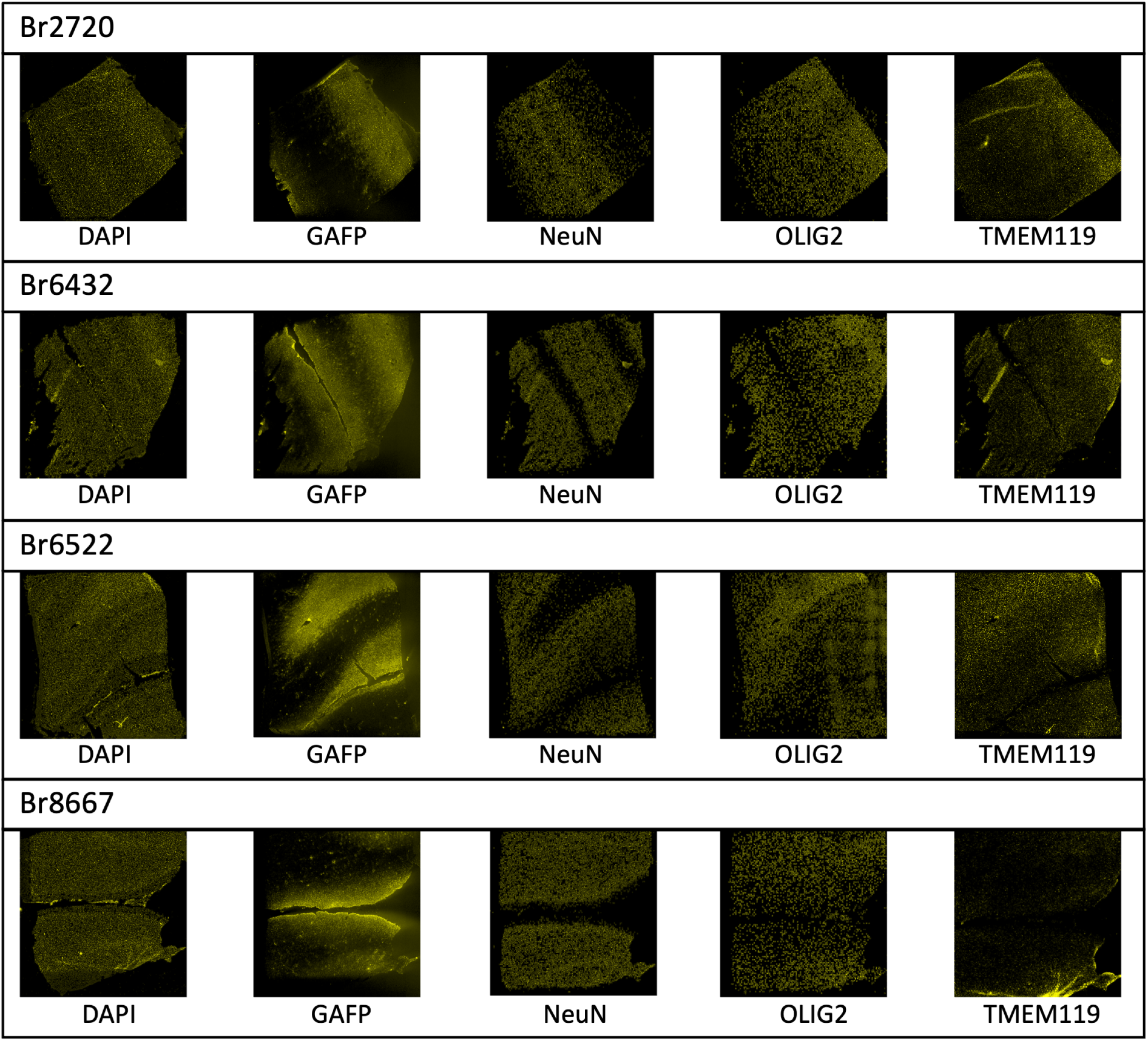
IF images from the Visium SPG human DLPFC samples. IF images of five cell-type channels (DAPI, GAFP, NeuN, OLIG2, and TMEM119) from four Visium SPG human DLPFC samples.

**Figure S4:**
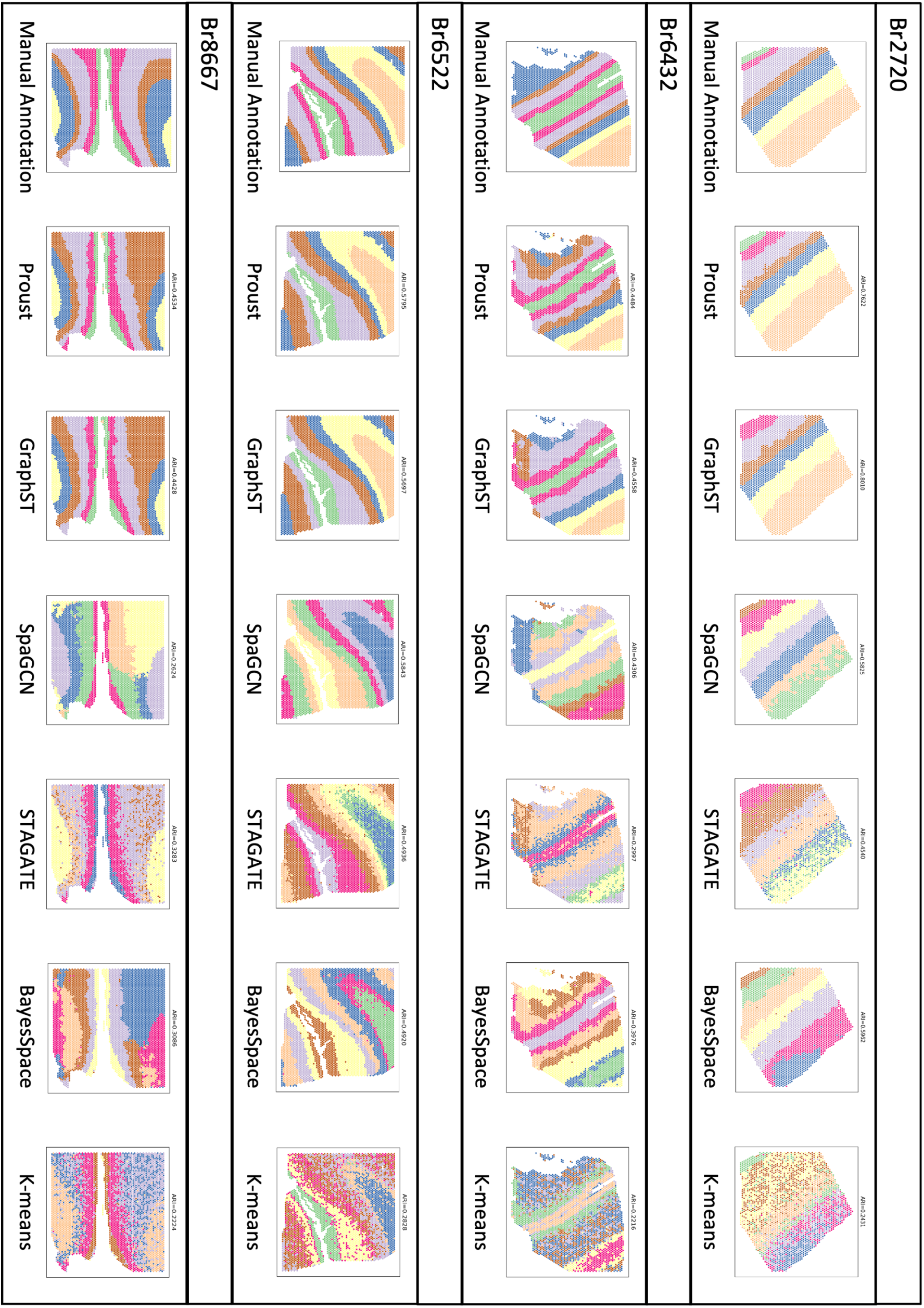
Manual annotations and clustering results from six methods on the Visium SPG human DLPFC samples. Manual annotations and clustering results from six methods on four Visium SPG human DLPFC samples.

**Figure S5:**
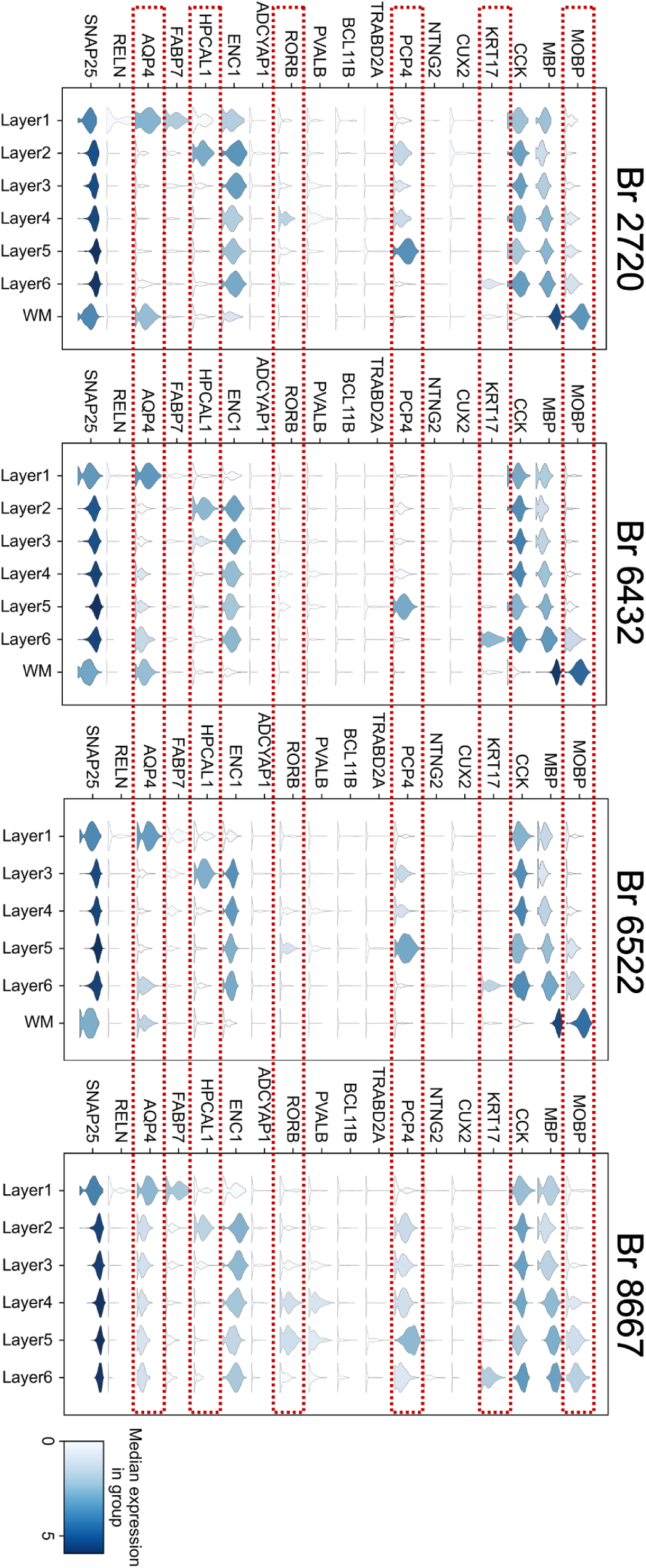
Stacked violin plots of marker genes across annotated clusters generated by Proust for the Visium SPG human DLPFC samples. Stacked violin plots of marker genes across annotated clusters generated by Proust for four Visium SPG human DLPFC samples. Selected marker genes for each layer are boxed.

**Figure S6:**
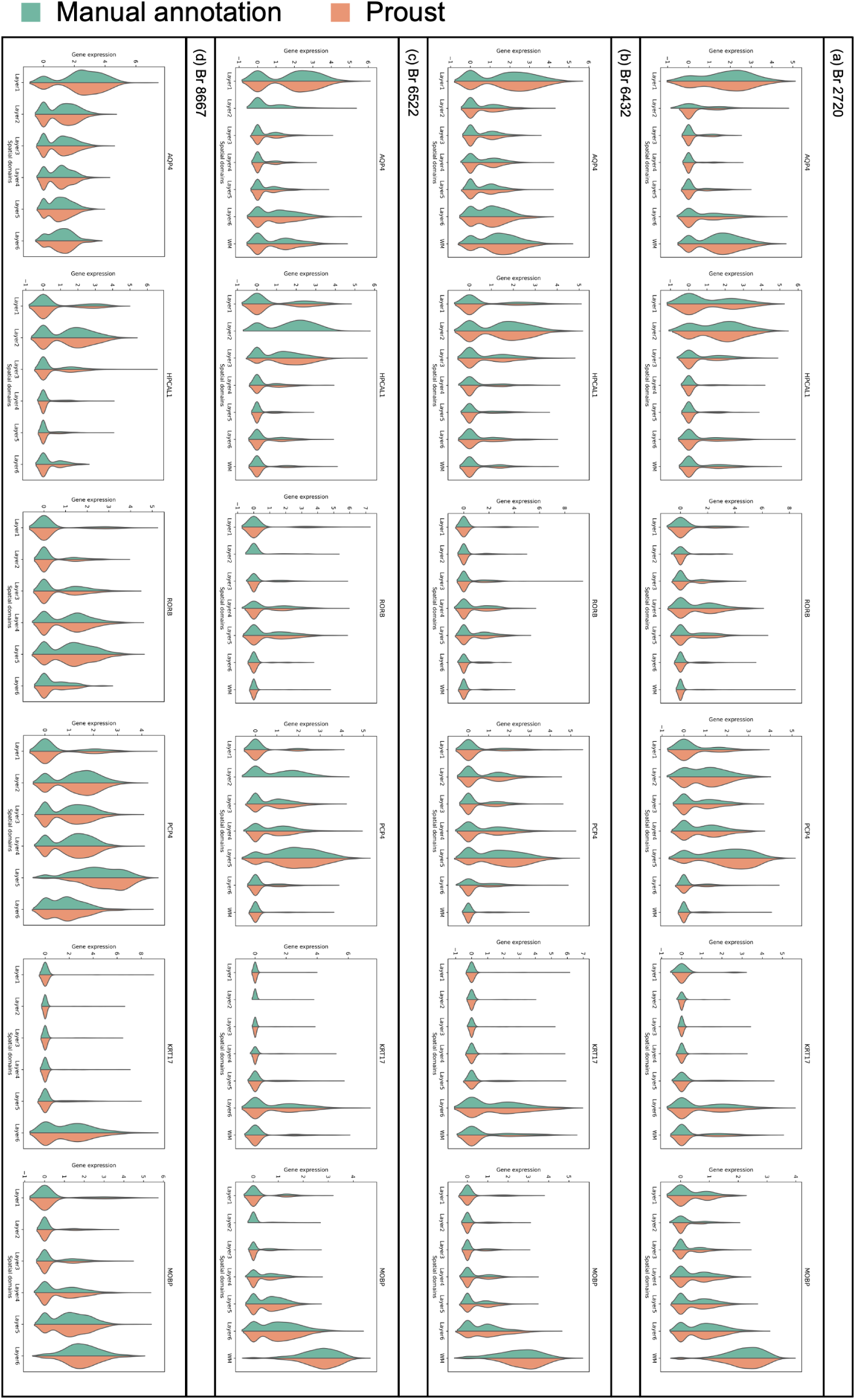
Violin plots of marker genes within clusters identified by Proust and manual annotations of the Visium SPG human DLPFC samples. Violin plots to compare marker gene distributions within clusters identified by Proust and manual annotations of four Visium SPG human DLPFC samples.

**Figure S7:**
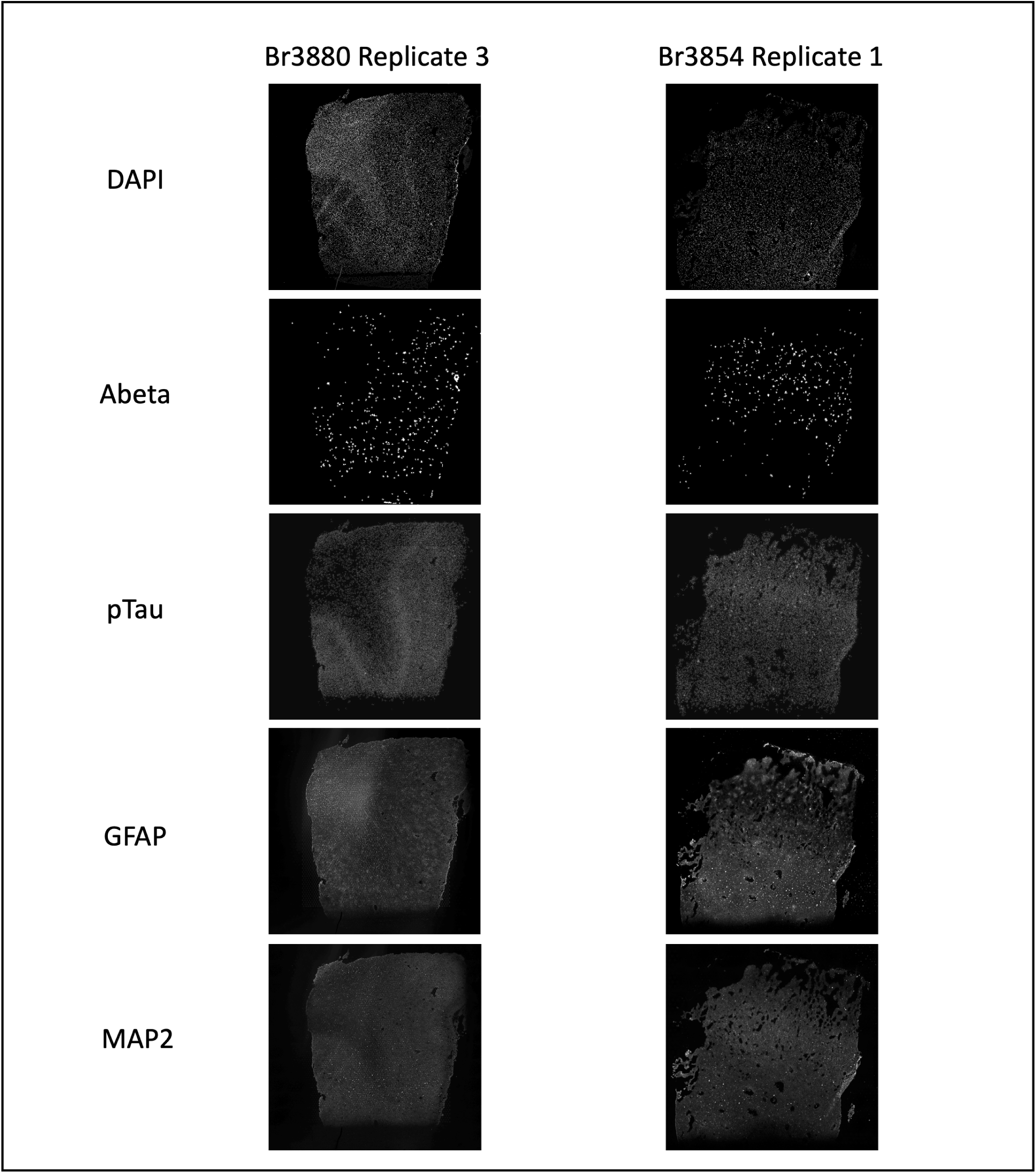
IF images from selected Visium SPG human inferior temporal cortex samples. IF images of five protein channels (DAPI, Abeta, pTau, GFAP, and MAP2) from selected Visium SPG human inferior temporal cortex samples.

**Figure S8:**
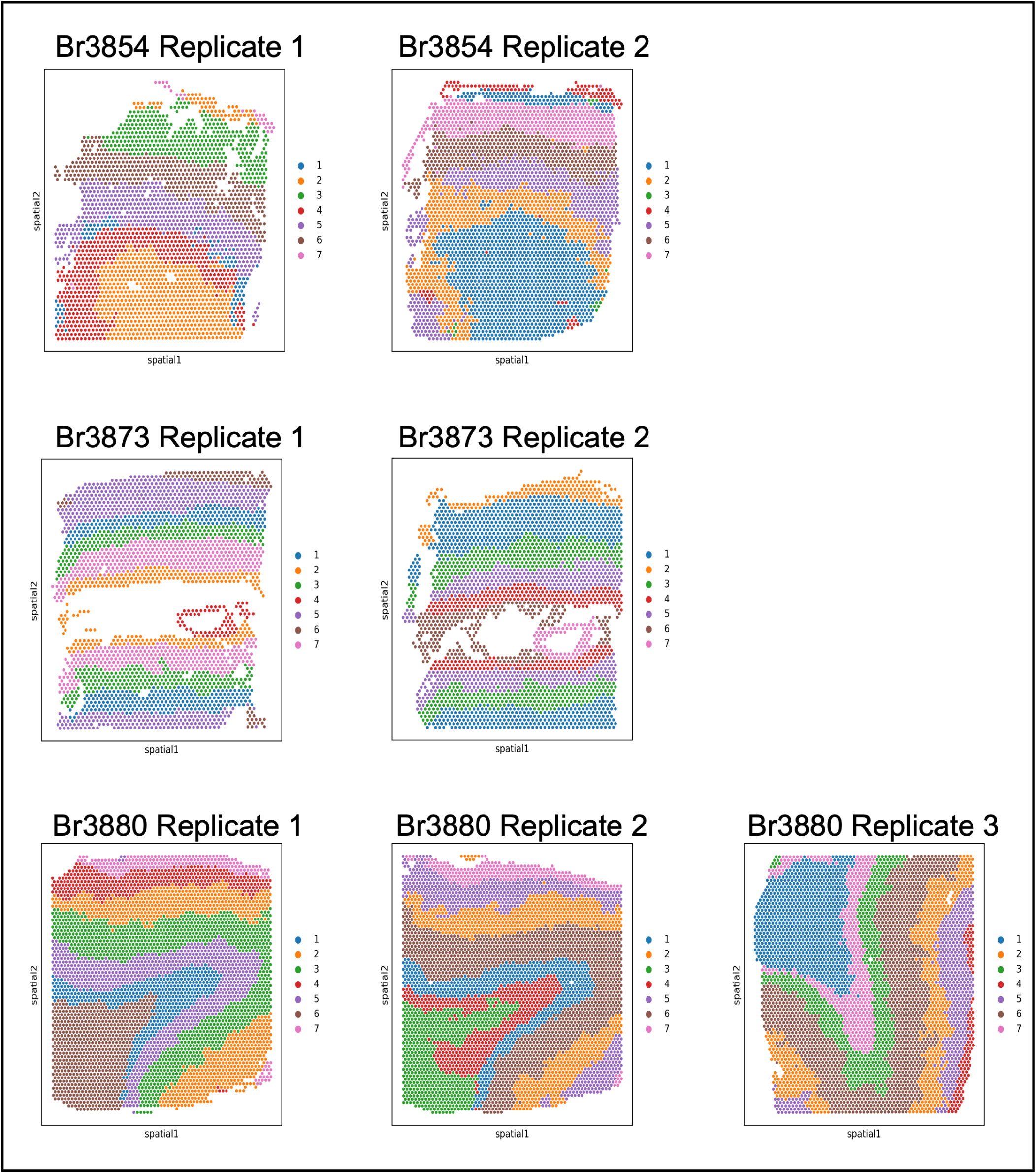
Proust clustering results of the Visium SPG human inferior temporal cortex samples, using five protein channels (DAPI, Abeta, pTau, MAP2, and GFAP), top 30 PCs from reconstructed gene expression, top 5 PCs from reconstructed extracted image features, and k = 7 clusters in Proust. Proust clustering results of seven Visium SPG human inferior temporal cortex samples, using five protein channels (DAPI, Abeta, pTau, MAP2, and GFAP), top 30 PCs from reconstructed gene expression, top 5 PCs from reconstructed extracted image features, and *k* = 7 clusters in Proust.

**Figure S9:**
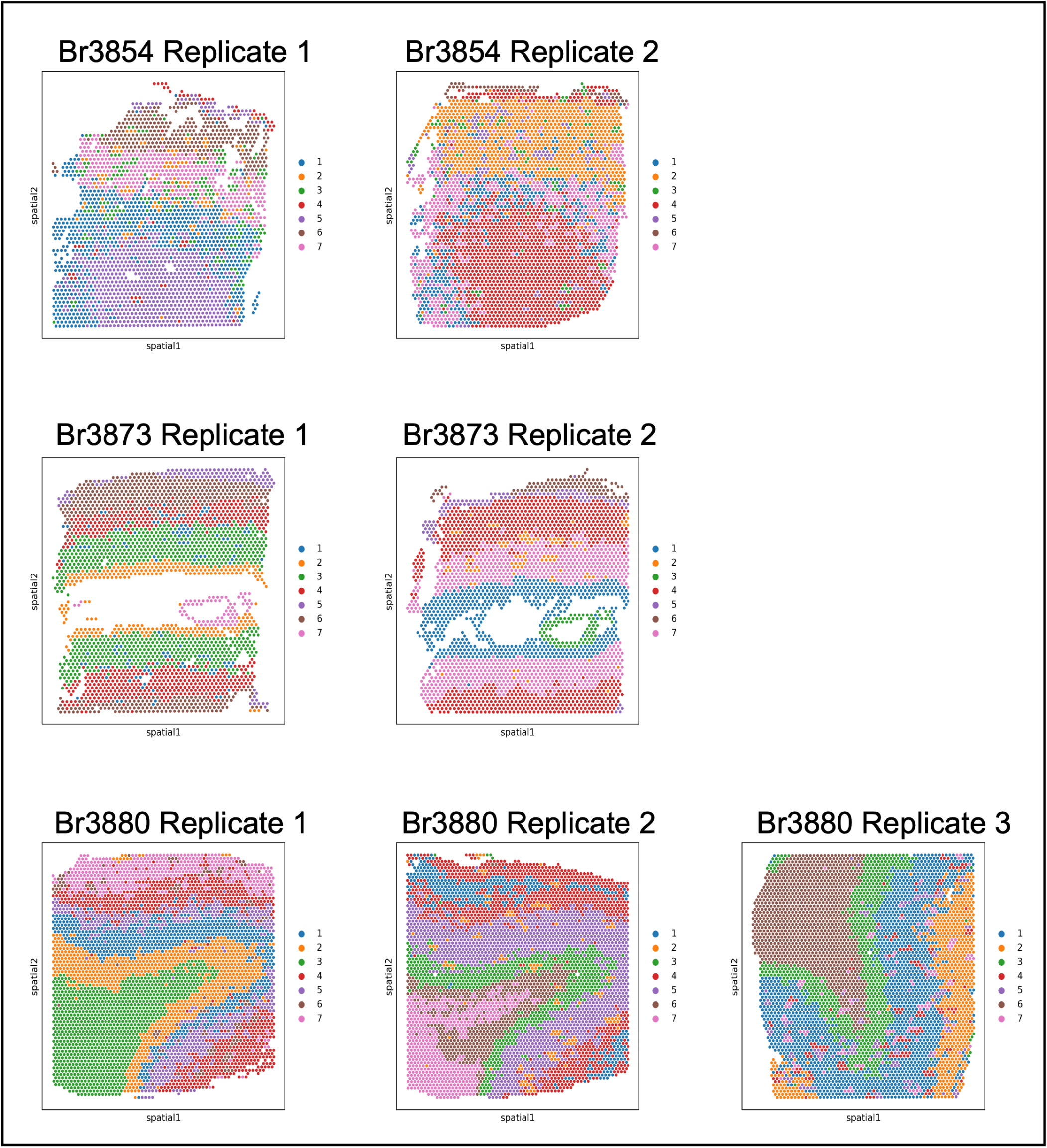
Proust clustering results of the Visium SPG human inferior temporal cortex samples, using two protein channels (Abeta and pTau), top 10 PCs from reconstructed gene expression, top 10 PCs from reconstructed extracted image features, and k = 7 clusters in Proust. Proust clustering results of seven Visium SPG human inferior temporal cortex samples, using two protein channels (Abeta and pTau), top 10 PCs from reconstructed gene expression, top 10 PCs from reconstructed extracted image features, and *k* = 7 clusters in Proust.

**Figure S10:**
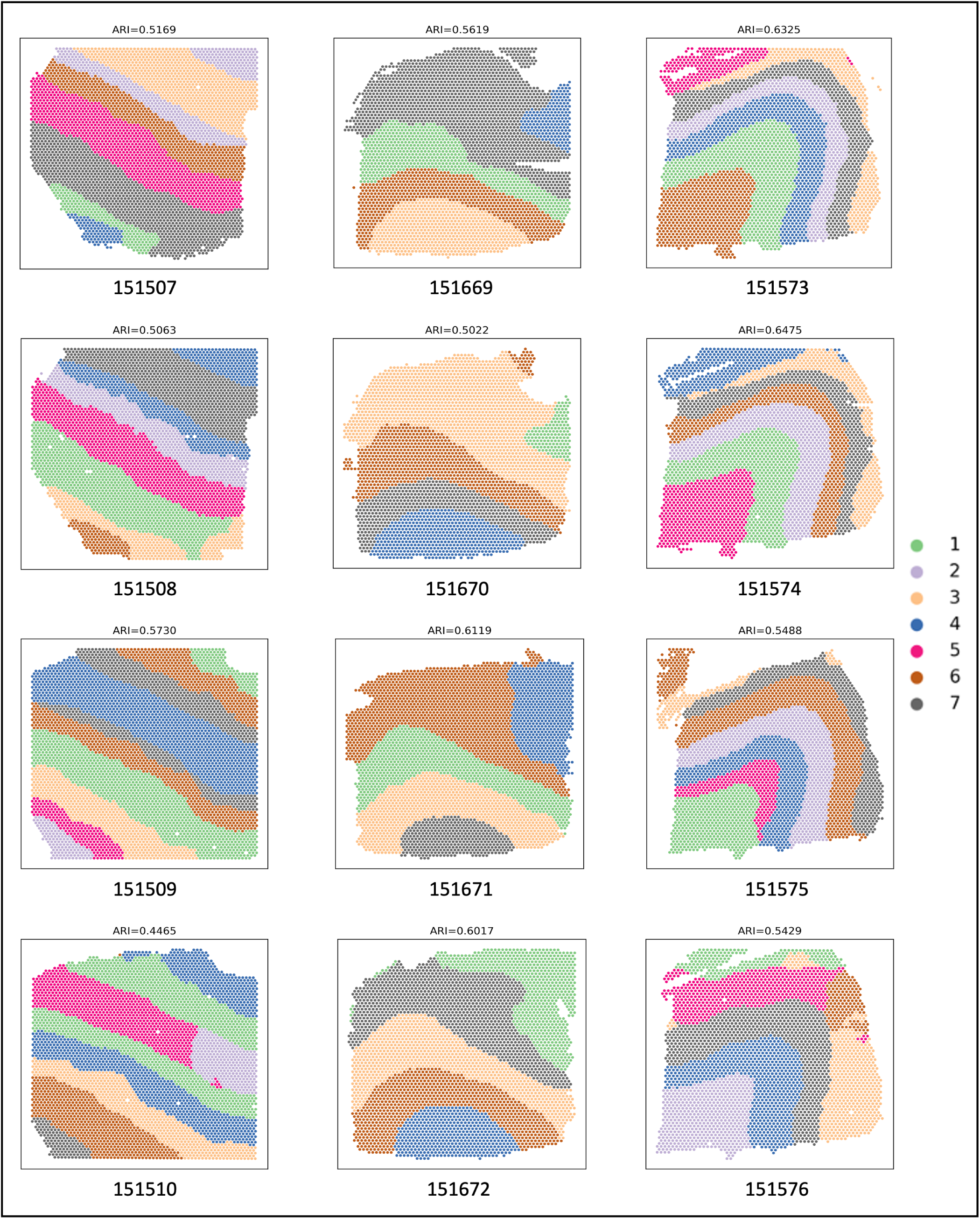
Proust clustering results of the Visium human DLPFC samples with H&E images. Proust clustering results of 12 Visium human DLPFC samples that contain H&E images.

